# In-vitro synergistic antibacterial effect of atorvastatin and ampicillin against resistant *Staphylococcus spp* and *E.coli* isolated from bovine mastitis

**DOI:** 10.1101/695817

**Authors:** Sriraam Sankar, Ramasamy Thangamalai, Sriram Padmanaban, Porteen Kannan, M R Srinivasan, C S Arunaman

## Abstract

The colossal rise in antimicrobial resistance has led to treatment failures and so mastitis has become cumbersome to treat. The objective of this study was to evaluate the antibacterial effect of non-antibiotic drug, atorvastatin in combination with antimicrobial, ampicillin against two commonly isolated bacterial species *Staphylococcus spp* and *E. coli* from bovine mastitis. Milk samples were collected from mastitis cows, visiting Veterinary Clinical Complex. Bacterial isolation was performed using Eosin Methylene Blue (EMB) agar and Mannitol Salt Agar (MSA), followed by characterization and identification by biochemical tests and gram staining. Genotypic confirmation was done by Polymerase Chain Reaction (PCR) with subsequent screening for resistant genes-*mec A*, bla_TEM_. Antibiotic Sensitivity Test (ABST) of the isolates against 12 different antimicrobials, atorvastatin only, and combination of atorvastatin with ampicillin were performed using Kirby-Bauer disc diffusion method. Minimum Inhibitory Concentration (MIC) of ampicillin alone and ampicillin in combination with atorvastatin were determined by modified microdilution method. *Staphylococcus spp* (77.5%) and *E.coli* (35%) were the two major pathogens isolated in the current study and multi-drug resistance was observed. Among the antimicrobials, the ampicillin showed 100% resistance against Staphylococcus spp and 85.71% resistance against E. coli. Atorvastatin did not display antibacterial effect as a sole agent but displayed synergistic antibacterial activity with ampicillin. There was an average increase in Minimum Inhibitory Concentration of ampicillin for *E.coli* and Staphylococcus spp isolates and atorvastatin decreased the Minimum Inhibitory Concentration of ampicillin in combination. The ampicillin shows more resistance against both *Staphylococcus spp* and *E.coli*, while atorvastatin improves the effect of ampicillin in-vitro. So, atorvastatin may be combined with ampicillin for the treatment of Gram-positive and Gram-negative infections. However, further studies are required to ascertain the exact mechanism of action of atorvastatin with respect to their antibacterial effect for them to be redeployed as an antimicrobial drug in the future.

## Introduction

Mastitis is the inflammation of the udder, caused by various pathogens like bacteria, fungi, protozoa, virus, and algae in milk producing animals. Mastitis is the costliest disease affecting dairy animals worldwide. In India, bovine mastitis leads to stupendous economic loss to the farmers due to reduced milk yield, poor quality of milk produced with subsequent food safety and public health threat. Antimicrobial resistance is a global health threat and it is starkly lofty in India. The nexus between antimicrobial resistance and mastitis pathogens is a renowned shackle to the dairy sector. In India, the total economic losses due to mastitis were calculated to be 7165.51 rupees. [1] *Staphylococcus spp* and *E. coli* are the two major mastitis pathogens in India and *Staphylococcus spp* is more prevalent than other mastitis pathogens in India. [2] Addition of amino group to the benzylpenicillin molecule yielded a drug with broadened spectrum of activity called “Ampicillin”. But, an increasing number of coliforms have become resistant to ampicillin. [3] With the emergence of “Superbugs” and diminished returns from the discovery of antibiotics, many pharmaceutical industries have refrained from discovery and development of new antibiotics. This gap can be effectively bridged by redeployment of available drugs as antibacterial agents. Statins is one such class of drugs with promising pleiotropic effects including antibacterial and anti-inflammatory properties [4]. In this view, this study was performed to evaluate the synergistic activity of atorvastatin with ampicillin against *Staphylococcus spp* and *E. coli.*

## Results

The present study reports that out of 100 milk samples collected from cows with mastitis, 60 samples (60%) showed growth on MSA and/or EMB agar. Remaining 40 samples showed growth in Nutrient Agar but not in MSA or EMB agar. Out of 60 samples, 40 samples were produced golden yellow colonies to pale white colonies in MSA agar (66.67%) and other 20 samples streaked onto EMB agar were produced metallic green sheen to purple coloured colonies (33.33%). Among the 60 samples, 6 samples (10%) produced colonies in both MSA and EMB agar, indicative of polymicrobial or mixed infection.

In this study, the Gram Staining of the presumptive isolates revealed that 60 samples showed growth on selective media. In that 40 isolates were observed to be Gram-Positive cocci (66.66%) and 20 isolates were observed to be Gram-Negative bacilli (33.33%).

In this study, the isolates were characterized as *Staphylococcus spp* (29/40) and *E.coli* (7/20) by using commercial biochemical test kits.

Presumptive isolates were subjected to PCR to confirm the presence of *S. aureus* and *E. coli* genotype. For confirmation of *Staphylococci spp*, *tuf* gene was targeted (Fig 1) To confirm the presence of *S. aureus, nuc* gene was targeted (Fig 2). Presence of *E. coli* was confirmed by targeting and amplifying *usp A* gene (Fig 3). Out of 40 isolates screened, 31 isolates were positive for *tuf* gene (77.5%). Among the 31 isolates confirmed to be *Staphylococci spp*, 13 isolates were confirmed as *S. aureus* (41.94%) due to the presence of *nuc* gene. Out of 20 isolates screened, 7 isolates were positive for *usp A* gene and therefore confirmed to be *E. coli* (35%).

**Fig 1.**
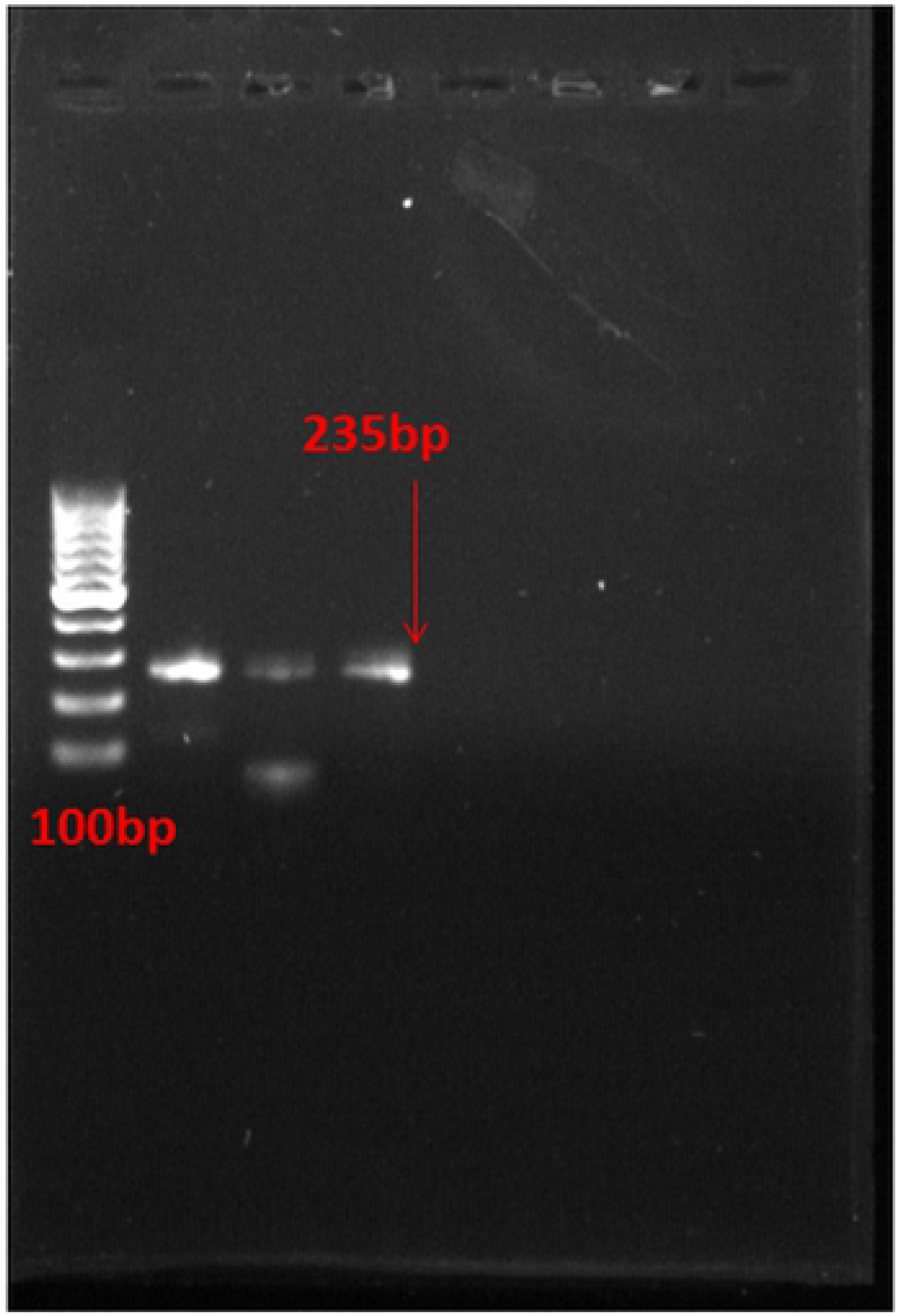
Presence of *tuf* gene

**Fig 2.**
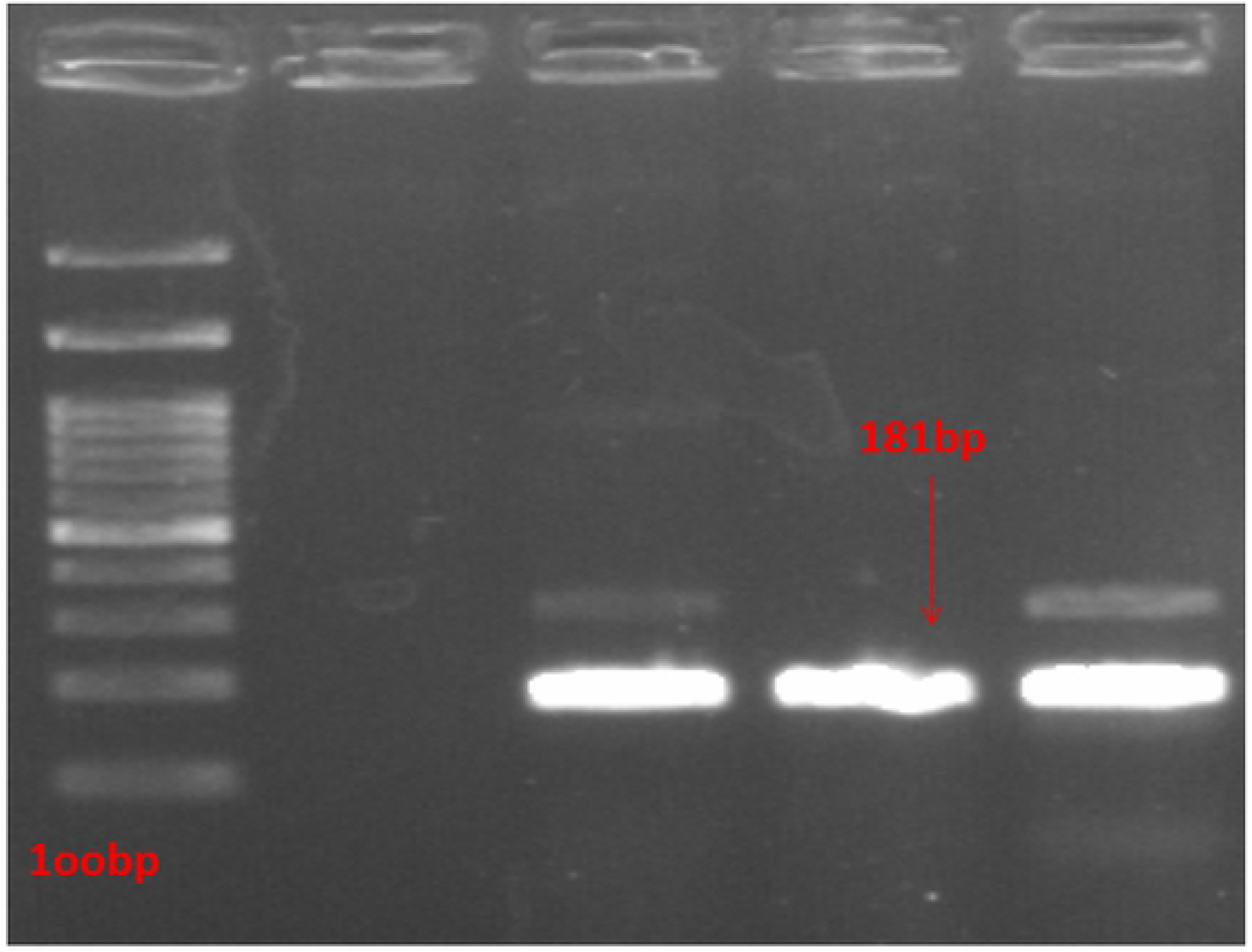
Presence of *nuc* gene

**Fig 3.**
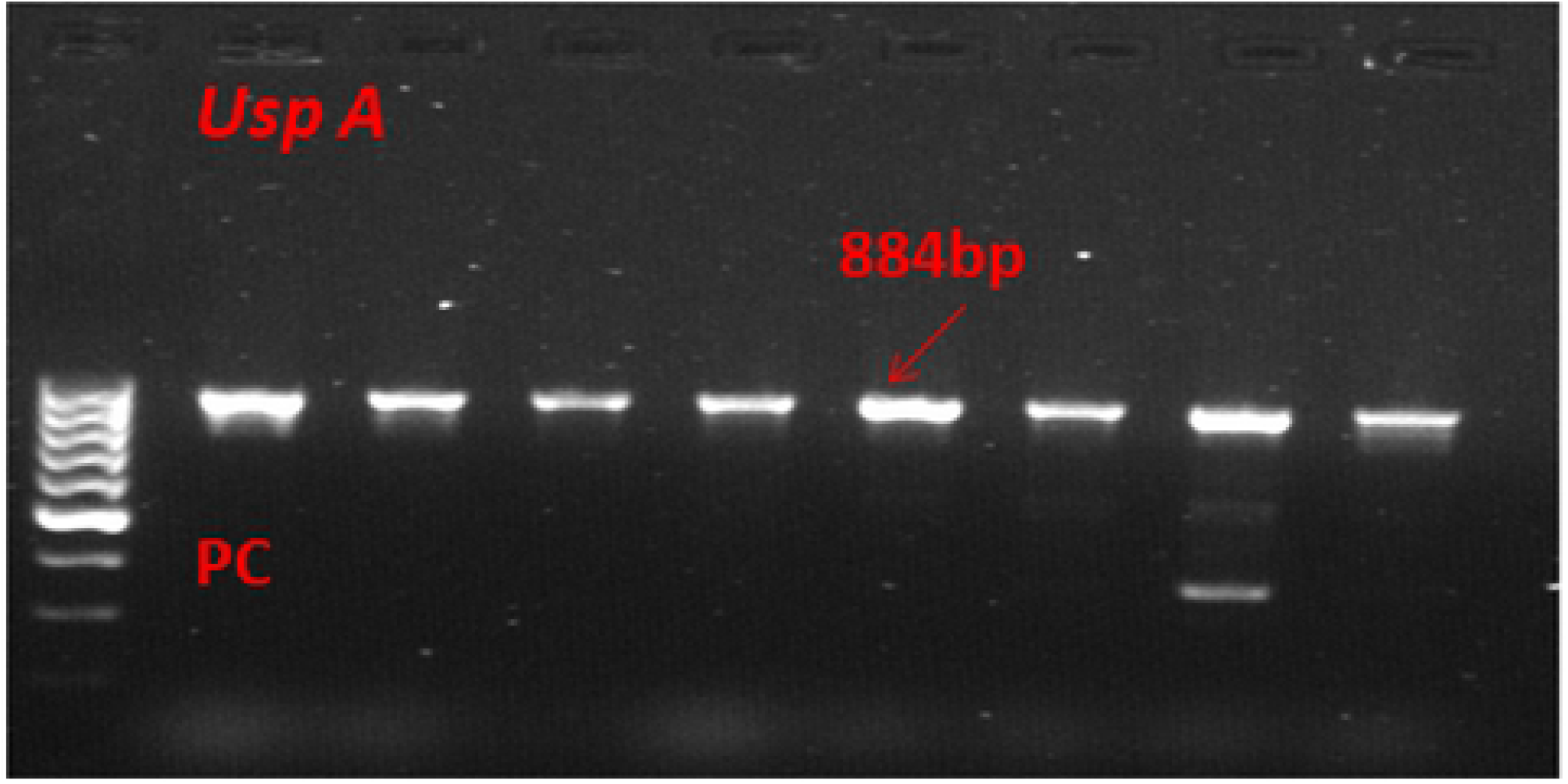
Presence of *usp A* gene

Out of 13 *S. aureus* isolates, 5 isolates (38.46%) were found to carry *mec A* gene (Fig 4), responsible for methicillin resistance and were identified as Methicillin-Resistant *Staphylococcus aureus* (MRSA). In the remaining 18 *Staphylococcus spp* isolates, 3 isolates (16.67%) were found to carry *mec A* gene, responsible for methicillin resistance. Out of 31 *Staphylococcus spp* isolates, 5 isolates (16.13%) were found to have bla_TEM_ gene (Fig 5) and out of 7 *E.coli* isolates screened, only one (14.29%) was found to be carrying bla_TEM_ gene, responsible for imparting ampicillin resistance.

**Fig 4.**
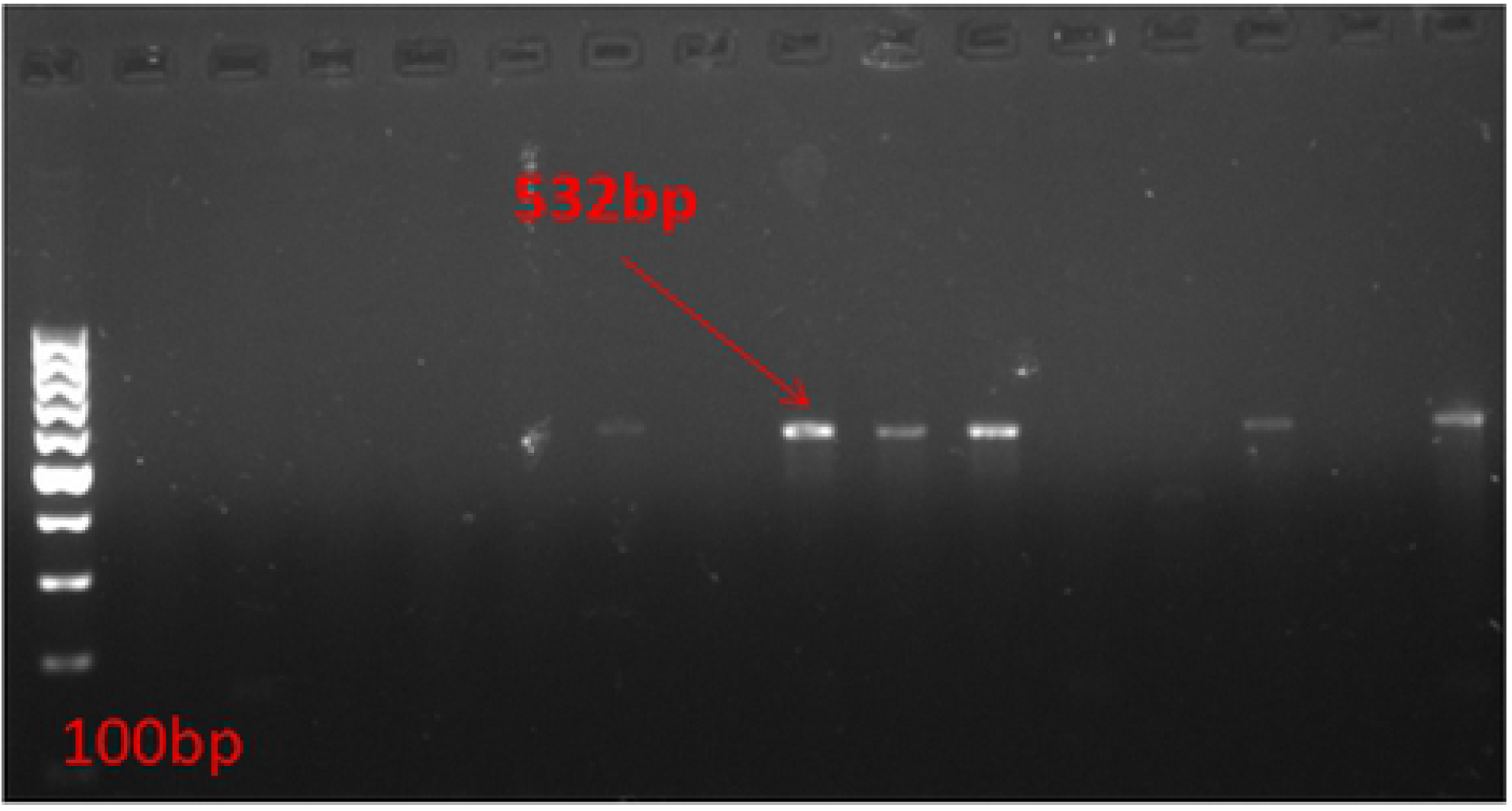
presence of *mec A* gene

**Fig 5.**
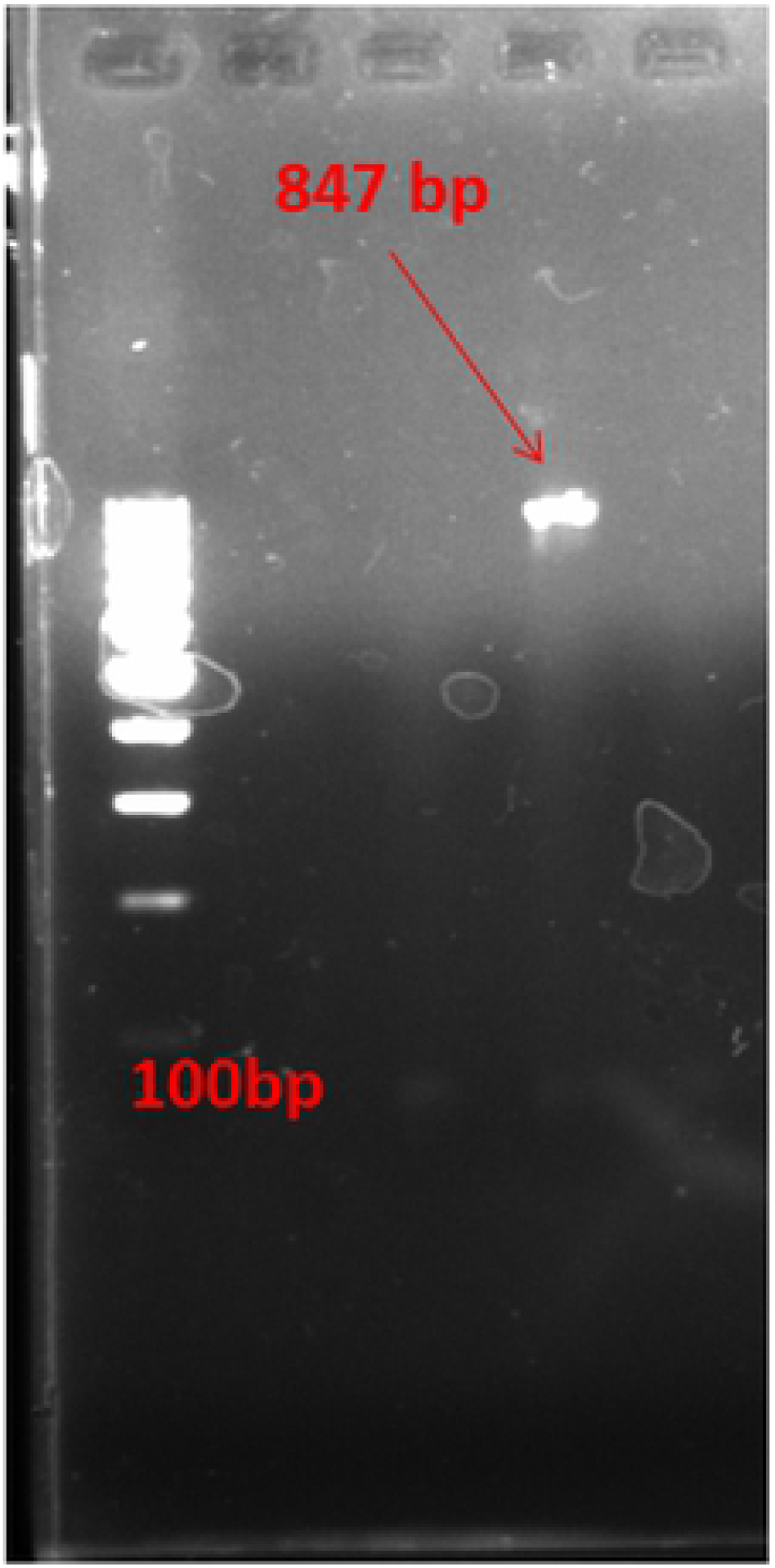
Presence of bla_TEM_ gene

The susceptibility test showed that in case of *Staphylococcus spp*, 80.65% samples were found to be resistant to tetracycline, gentamicin, vancomycin and amikacin. 58.06% samples were found to be resistant to cephotaxime, 61.29% samples were found to be resistant to ceftriaxone, 74.19% samples were found to be resistant to co-trimoxazole, 51.61% samples were found to be resistant to enrofloxacin, 61.29% samples were found to be resistant to ciprofloxacin, 83.87% samples were found to be resistant to methicillin. Complete resistance (100%) was observed against penicillin and ampicillin, phenotypically.

The susceptibility test showed that in case of *E.coli*, 85.71% samples were found to be resistant to tetracycline, gentamicin, cephotaxime, ceftriaxone, co-trimoxazole, ampicillin, amikacin and ciprofloxacin. Against enrofloxacin, 75.43% samples showed resistance. Complete resistance (100%) was observed against penicillin, methicillin and vancomycin.

In this study, evaluation of antibacterial effect of atorvastatin revealed no significant antibacterial activity against both *Staphylococcus spp* and *E.coli* isolates in disc diffusion method. The atorvastatin and ampicillin combination showed increased zones of inhibition against both *Staphylococcus spp* and *E.coli* isolates.

The average increase in MIC of ampicillin for *E.coli* isolates was 22.62μg/ml (Fig 6). The recommended CLSI (2016) MIC breakpoint for ampicillin against E.coli is ≤ 8 μg/ml. The average increase in MIC of ampicillin for *Staphylococcus spp* isolates was 17.20 whereas the recommended MIC for ampicillin against *Staphylococcus spp* by CLSI (2016) is ≤4 μg/ml. The average decrease in MIC of ampicillin in combination with atorvastatin against *Staphylococcus spp* and *E.coli* was 2.49μg/ml and 2.61μg/ml, respectively. The results are depicted in the Table 1.

**Fig 6.**
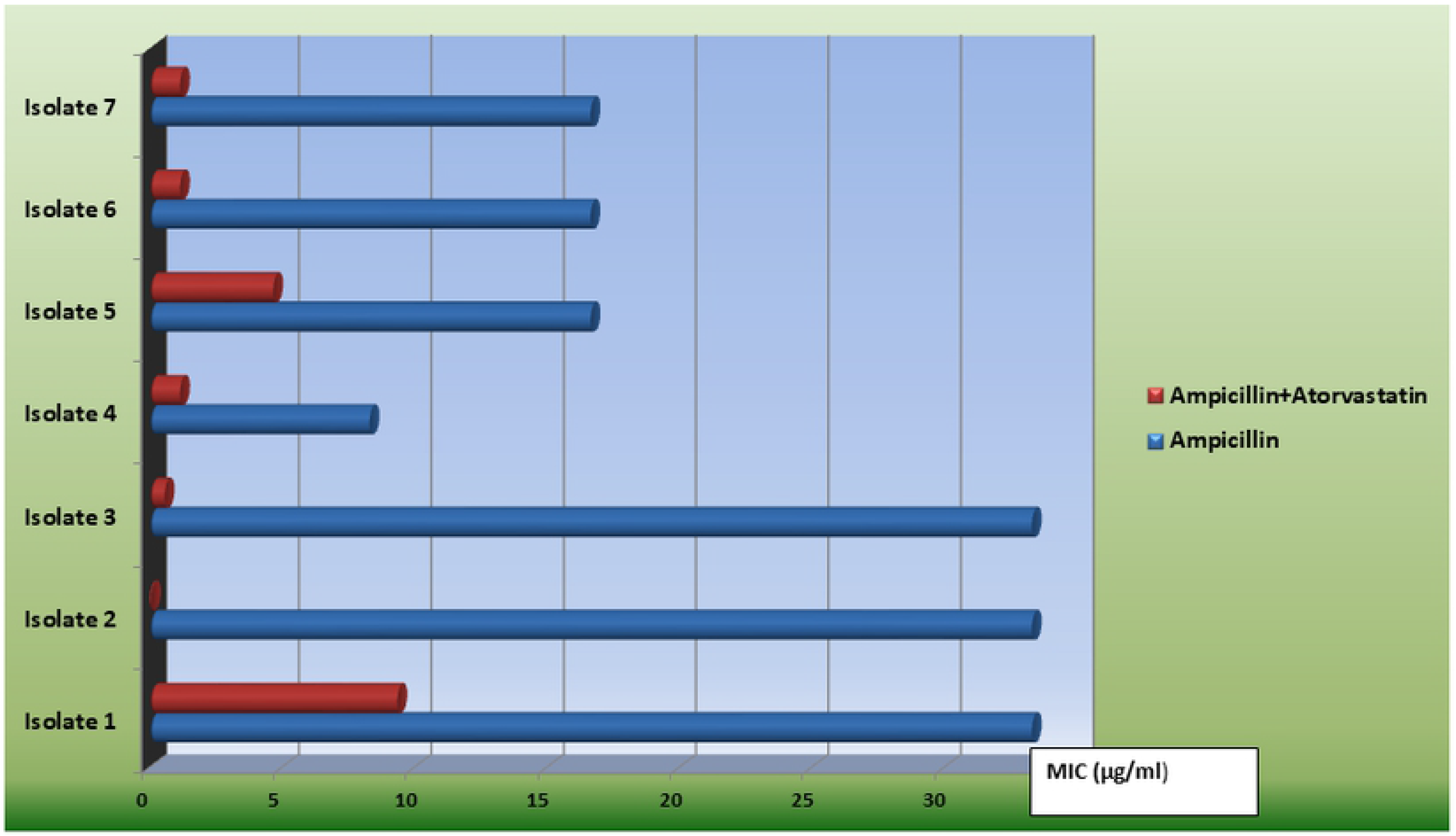
Comparison of MIC for *E. coli* isolates

**Table 1.** Comparison of MIC for ampicillin and ampicillin with atorvastatin combination

| Group | MIC( Mean ± SE ) | |
| --- | --- | --- |
|  | E.coli | Staphylococcus spp |
| Ampicillin | 22.62 ± 3.95<sup>b</sup> | 17.20 ± 1.44<sup>b</sup> |
| Ampicillin + Atorvastatin | 2.61 ± 1.26<sup>a</sup> | 2.49 ± 0.49<sup>a</sup> |

Mean ± SE bearing different superscripts differ significantly at 1% level of significance. (p<0.001)

## Discussion

In the present study, *Staphylococcus spp* was the predominant bacteria isolated from the mastitis milk samples, followed by *E. coli*. This finding is in complete agreement with many other previous studies, including the studies by Jeykumar *et al.* (2013) [5] and Padhy *et al.* (2014) [6], which also reported that *Staphylococcus spp*, followed by *E. coli* were the two important pathogens isolated from mastitis milk samples. A study by Hazlett *et al*. (1984) [7] also reported mixed bacterial infection as a cause of bovine mastitis in 22.2% of the cases encountered in the study and also reported that mixed infections were fairly common with *E. coli*. This finding is also line with the finding of the current study.

In the current study, Out of 40 presumptive isolates screened, 31 isolates were positive for *tuf* gene (77.5%). Among the 31 isolates confirmed to be *Staphylococci spp*, 13 isolates were confirmed as *S. aureus* (41.94%) due to the presence of *nuc* gene. This finding correlates with the finding of the study conducted by Xu *et al.* (2015) [8]. Out of 20 presumptive isolates screened, 7 isolates were positive for *usp A* gene and therefore confirmed to be *E. coli* (35%). and a similar finding was reported by Mishra *et al.* (2017) [9]. The pattern of phenotypic resistance for the tested antimicrobial drugs in this study was similar to the pattern reported in various other studies by [10], [11], [12] and [13].

None of the isolates was found to be sensitive to atorvastatin when used as a sole agent. The finding that atorvastatin did not exhibit antibacterial effect on its own is in stark contrast with many other previous studies [14], [15] but this finding is in complete agreement with the study conducted by Manalo *et al*. (2017) [16], which also reported that atorvastatin did not display any antibacterial activity, when used as a sole agent.

In the present study, it was estimated that the average increase in MIC of ampicillin for *E.coli* isolates was 22.62μg/ml, while the average increase in MIC of ampicillin for *Staphylococcus spp* isolates was 17.20μg/ml. The combination of ampicillin with atorvastatin exhibited synergistic antibacterial activity against both *Staphylococcus spp* and *E.coli*, with an average decrease of MIC to 2.49μg/ml and 2.61μg/ml for *Staphylococcus spp* and *E.coli*, respectively. This result of the current study is consistent with the finding by Manalo *et al.* (2017).

## Methods

Milk samples from suspected cases of bovine mastitis were aseptically collected and stored at 4°C, till they were processed further. Selective isolation of *Staphylococcus spp, E. coli* was performed using Mannitol Salt Agar (MSA) and Eosin Methylene Blue agar (EMB). The individual colonies obtained from selective media were stored as glycerol stocks at −20°C and they were used for all further testing procedures. Biochemical kits were used for characterization of the isolates (HiMedia^TM^). PCR was performed for genotypic characterization of the isolates [17] including *tuf* gene for *Staphylococcus spp* [18], *usp A* gene for *E.* coli [19], *nuc* gene for *S. aureus* [20]. PCR was carried out for detection of resistance gene bla_TEM_ for ampicillin resistance [21]. The amplified PCR products were separated using electrophoresis on a 1.2% agarose gel with ethidium bromide (10% of the total volume) in 1X SBB buffer. The separated PCR products in the gel were visualized on Mega Capt Gel Doc. A 100bp DNA ladder (Thermoscientific^TM^) was employed to determine the size of the PCR gene products.

**Table 2.**
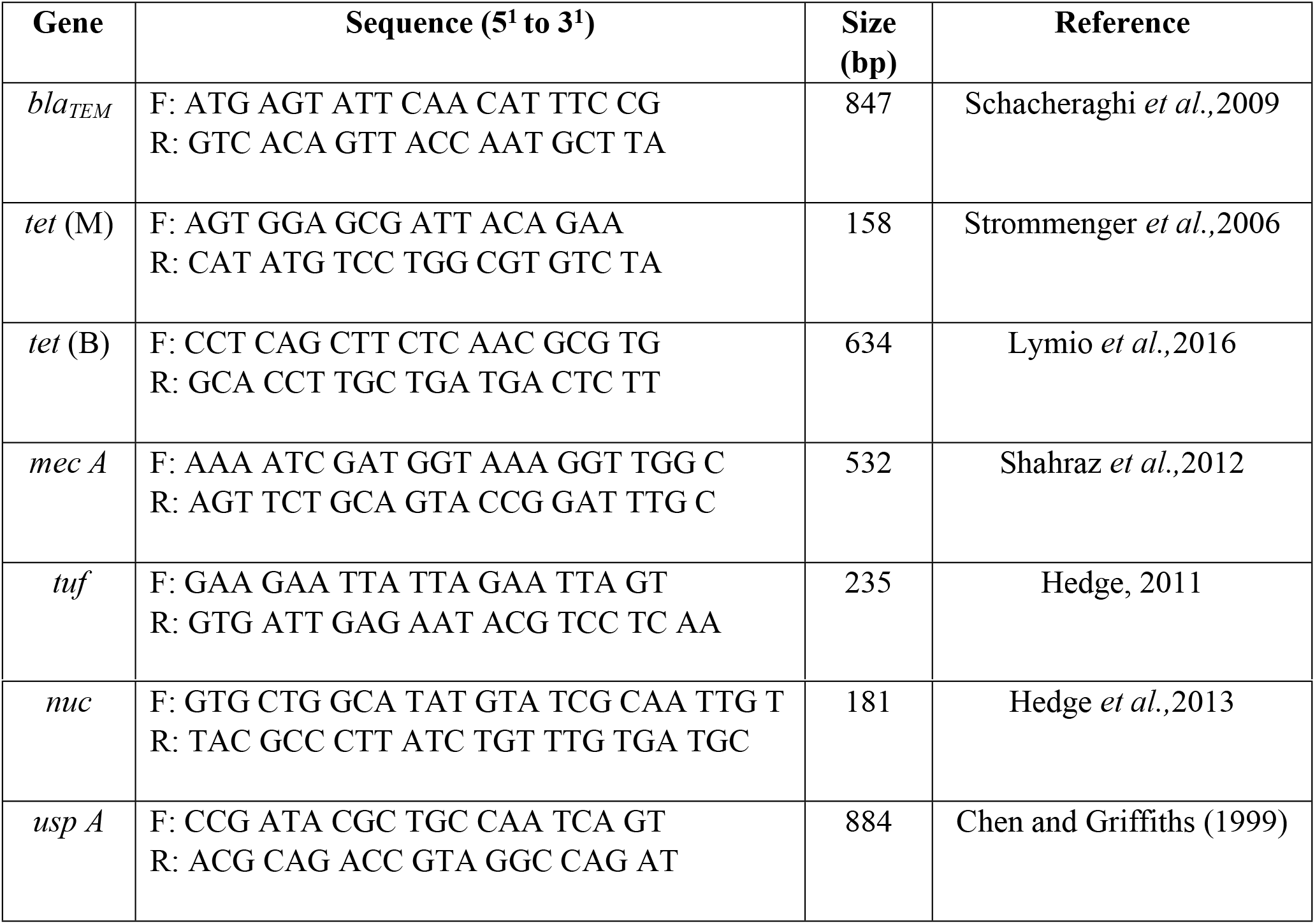
List of primers used in this study

**Table 3.**
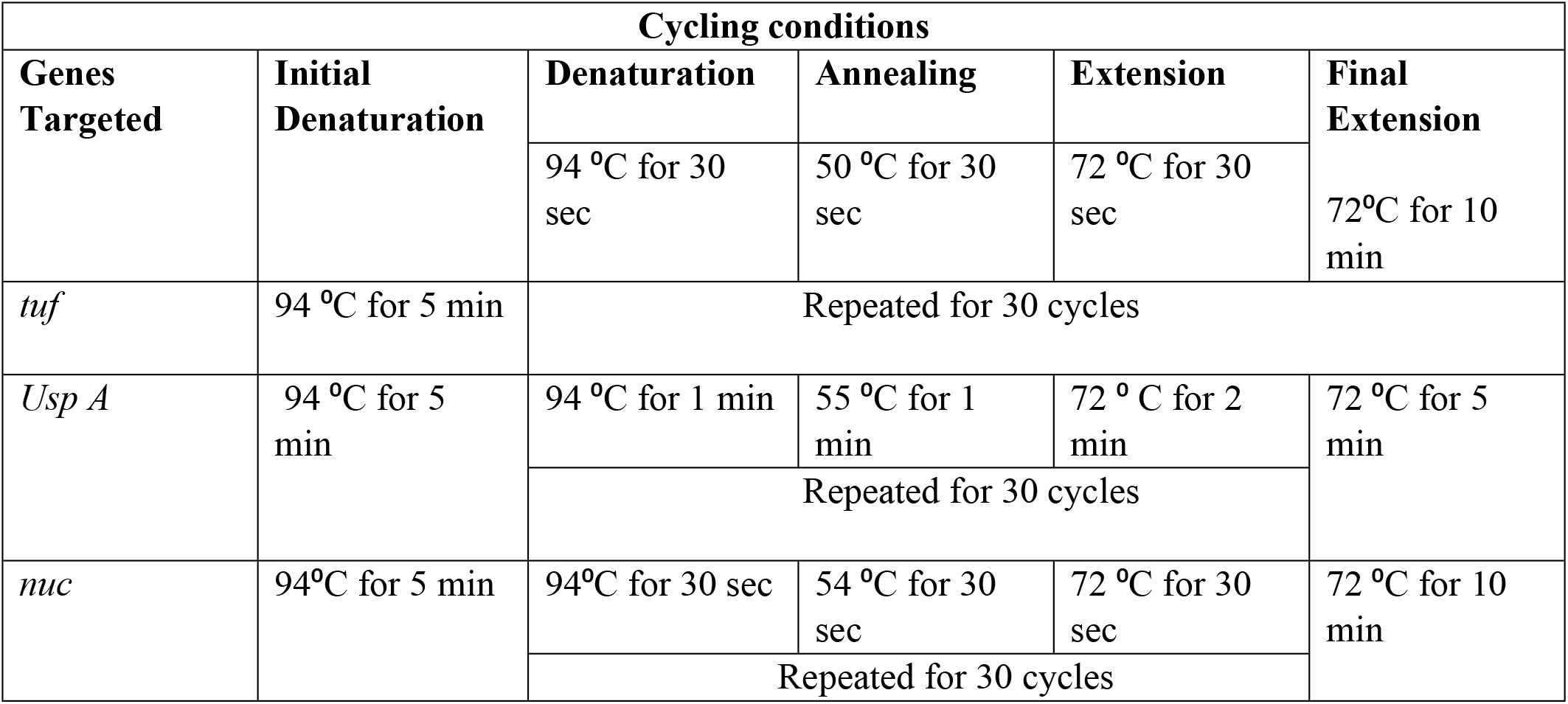
PCR cycling conditions for identification genes

**Table 4.** PCR cycling conditions for resistance genes

| Cycling conditions |  |  |  |  |  |
| --- | --- | --- | --- | --- | --- |
| Genes Targeted | Initial Denaturation | Denaturation | Amealing | Extension | Final Extension |
|  |  | 94 °C for 1 min | 55 °C for 1 min | 72 °C for 2 min |  |
| *mec A* | 94 ° C for 5 min | Repeated for 35 cycles |  |  | 72 °C for 5 min |
| *tet* (B) | 95 ° C for 5 min | 95 °C for 30 sec | 55 °C for 30 sec | 72 °C for 30 sec | 72 °C for 2 min |
|  |  | Repeated for 30 cycles |  |  |  |
| *tet* (M) | 94 ° C for 5 min | 94 ° C for 1 min | 50 ° C for 1 min | 72 ° C for 90 sec | 72 °C for 5 min |
|  |  | Repeated for 30 cycles |  |  |  |
| *bla<sub>TEM</sub>* | 96 °C for 5 min | 96 °C for 1 min | 58 °C for 1 min | 72 °C for 1 min | 72 °C for 10 min |
|  |  | Repeated for 35 cycles |  |  |  |

ABST was performed using modified Kirby-Bauer disc diffusion method. An overnight inoculum of culture was diluted to 0.5 McFarland standard and used for ABST. All the antibiotic discs were purchased from HiMedia^TM^. For synergy evaluation, 1mg of atorvastatin was dissolved in 1ml of Dimethyl Sulfoxide (DMSO) and 8 μl of the same solution was added to the ampicillin discs. The discs were allowed to be set for 20 minutes at room temperature and then incubated at 35-37°C for 24 hours. A sterile disc inoculated with same volume of DMSO served as control [22].

MIC (Minimum Inhibitory Concentration) was evaluated using modified microdilution method using resazurin indicator method. A 96 well plate was used for testing of samples and resulting colour change was noted and the concentration of the drug in the well was calculated to find MIC [23].

## Acknowledgements

We would like to acknowledge the help of Dr. Balan, Assistant professor, Department of Statistics, Madras veterinary College, Chennai 600007.

---

S1 Text. **Zones of inhibition (mm) for *Staphylococcus spp* isolates of various antimicrobial drugs.**

S2 Text. **Zones of inhibition (mm) for *E. coli* isolates of various antimicrobial drugs.**

